# Identification of sex-biased miRNA markers informative of heat-past events

**DOI:** 10.1101/2024.01.24.577069

**Authors:** Tosca A. van Gelderen, Jerome Montfort, José Antonio Álvarez-Dios, Francesc Piferrer, Julien Bobe, Laia Ribas

## Abstract

Elevated temperatures during early developmental stages play a pivotal role in the fate of the final sexual phenotype of fish populations, particularly towards male-skewed sex ratios. This is the case with European sea bass (*Dicentrarchus labrax*), one of the most important European species in the aquaculture industry. To unveil informative markers of the past thermal events, we investigated changes in the miRNome within the gonads of this species. Consequently, we exposed European sea bass to elevated temperatures during early development and conducted miRNA-sequencing analysis in the ovaries and testes one year post-heat treatment. The examination of miRNA expression levels identified three and twelve miRNAs in ovaries and testes, respectively, reflecting past thermal events. To assess the evolutionary conservation of these identified miRNAs in gonads, we cross-referenced our data with miRNome public information from ovaries and testes in nine additional fish species. This analysis uncovered 43 potential sex-biased markers present in at least five studied species along the evolutionary timeline. For instance, miR-155, miR-429, and miR-140 were female-skewed while miR-184, miR-499, and miR-135 were male-skewed. In addition, among these markers, eight conserved sex-skewed miRNAs proved informative regarding past thermal events in both the ovaries (e.g., miR-192-5p and miR-143-3p) and testes (miR-129-5p, miR-2187a-3p, miR-724-5p, miR-143-3p, miR-194a-5p and miR-223-3p). Notably, miR-223-3p and miR-194a-5p were conserved female-skewed, but showed upregulation in males exposed to high temperature. These miRNAs could serve as markers of heat-induced masculinization of genetic female-prone fish. The current research broadens the inventory of sex-specific miRNAs along evolution in fish and, elucidates thermosensitive miRNAs in the gonads. These findings hold promise as potential tools for predicting historical environmental events associated with masculinization due to high-temperature treatments in cultured species but also perhaps for natural populations exposed to a climate change scenario.

## Introduction

The European sea bass (*Dicentrarchus labrax*) is important for aquaculture. In 2016, aquaculture accounted for 96% of the total E. sea bass production in comparison with that obtained from fisheries (*FAO*, 2022; Vandeputte et al., 2019). To ensure successful growth and development in cultured fish, it is essential to maintain optimal rearing. For economic production efficiency, E. sea bass females are the desired sexual phenotype as they are favored due to their larger size, which can be up to 30% greater than their male counterparts (Ref.). However, farmed E. sea bass is known to produce populations consisting of 70-100% males, due to increased temperatures during early stages of development in the hatcheries (Piferrer et al., 2005; Saillant et al., 2002). Various studies have attempted to identify the ideal rearing temperature for E. sea bass larvae but it is known that temperatures exceeding 16 degrees Celsius (°C) during sex differentiation window (∼ 50 to 140 days post fertilization, dpf) lead to male-skewed populations (Navarro-Martín et al., 2009; Wang and Shen, 2018).

Unlike species with genotypic sex determination, some teleost fish, including the E. sea bass, exhibit polygenic sex determination (PSD) (Vandeputte et al., 2007). In PSD, environmental factors play a substantial role in shaping the final sexual phenotype of individuals. These external influences impact gene expression through various epigenetic mechanisms (Granada et al., 2018). The first description of epigenetics in sex regulation due to temperature in fish was given in E. sea bass more than a decade ago. DNA methylation of the gonadal aromatase cytochrome P450, family 19, subfamily A (*cyp19a1a*) promoter —a key gene to develop ovaries— was altered in adults previously reared at a high temperature during early development, favoring the masculinization of the final population (Navarro-Martín et al., 2011). Since then, endeavoring research has been deployed to better understand the connection between the epigenome and sexual phenotypes.

The function of miRNAs —small non-coding RNAs that play a vital role in post-transcriptional gene regulation— is to act as an epigenetic mechanism and serve as a bridge between the environment and the genome. In the last years, the importance of miRNAs in fish has flourished and thus research related to miRNA functions in cultured fish has grown exponentially. Many studies regarding temperature alterations and miRNA expression have been described in various species and tissues in fish. For example, after short-term and acute high or low temperature exposures (18 days), 29 miRNAs were differentially expressed (DE) in the common carp (*Cyprinus carpio*) liver (Sun et al., 2019). In the Antarctic notothenioid (*Trematomus bernacchii*), adapted to extreme climates of temperatures around −1°C, an acute heat stress was able to alter 12 miRNAs in the gills, mainly pertaining to cellular stress response (Vasadia et al., 2019). Long-term effects on the miRNome were shown in zebrafish (*Danio rerio*) (van Gelderen et al., 2022) and in Atlantic cod (Bizuayehu et al., 2015). Further, adult zebrafish exposed to high temperature for 21 days changed the miRNome with a consecutive recovery in which miRNA expressions returned to their original state (Ikert and Craig, 2020). In particular, in E. sea bass exposed at high temperature during different stages of early development, miRNA expression was assessed showing an alteration of heat-sensitive miRNAs in which the immune system and the reproduction system were involved (Papadaki et al., 2022).

miRNAs emerged during early evolution, i.e., in metazoans, and some miRNAs have been described as ancient miRNAs in bilaterians, such as miR-100, miR-125 and let-7 (Christodoulou et al., 2010). One of the particularities that makes miRNAs good markers for conserved developmental processes is their high conservation of both sequence and function along evolution (Niwa and Slack, 2007). Thus, ever since their initial discovery, miRNAs have been used as tools in extensive applications in human health diagnostics, primarily for tumor detection (Hamam et al., 2017). In the context of aquaculture, and since the first miRNA discovery in zebrafish (Lim et al., 2003), numerous studies have been undertaken to explore and characterize the miRNA repertoire in other teleost species. In fish, miRNA expression patterns were proposed as indicators to improve productivity by, for example, optimizing selection for breeding programs. This is the case of rainbow trout (*Onchorynchus mykiss*), in which circulating miRNAs in the blood were recently identified as a non-invasive approach to detail the metabolic and reproductive states (Cardona et al., 2021). In Nile tilapia miR-1, miR-206, and miR-133a abundance was suggested as markers to link genetic polymorphism data with miRNA targets (Huang et al., 2012). By using public databases, a set of miRNAs (i.e., miR-9-3p, miR-135c, miR-9-5p, miR-30b, miR-122 and miR-92a-3p) was suggested as a collection of potential biomarkers for cold tolerance in fish (Blödorn et al., 2021).

Notwithstanding the existence of many studies regarding the miRNA roles in the reproduction system, the functions of many miRNAs, particularly in the context of sex development, remain still enigmatic in fish. The elucidation of conserved miRNAs within the gonads of teleost fish species may provide valuable insights into the identification of markers relevant to sex development. In light of the relative lack of research on the E. sea bass species in this field, delving into the E. sea bass miRNome promises to offer valuable insights into the intricate relationship between the environment, miRNA expression patterns and the sexual phenotype. Thus, the aims of the present study were: 1) to enrich the knowledge of miRNA present in the E. sea bass gonads; 2) to identify sexual biomarkers in fish; 3) to discover heat recorders biomarkers of environmental past events like temperature. To achieve our goals, miRNA-sequencing strategies of E. sea bass gonads subjected to high temperatures were used. Furthermore, we undertook a comparative analysis of our E. sea bass miRNA data with obtained profiles of public gonadal miRNAs databases of nine different species (Desvignes et al., 2022). : zebrafish, three-spined stickleback (*Gasterosteus aculeatus*), striped catfish (*Pangasianodon hypophthalmus*), Japanese medaka (*Oryzias latipes*), black bullhead (*Ameiurus melas*), European perch (*Perca fluviatilis*) and eastern mudminnow (*Umbra pygmaea*) and two Holostean species: bowfin (*Amia calva)* and spotted gar (*Lepisosteus oculatus*) (Desvignes et al., 2018) were used in this study.

## Materials and methods

### Experimental design

The fish used in this study were siblings of those used in previous studies in which a significant male-skewed sex ratio (70.45% males to 29.55% females, *p-*value < 0.001) was observed after high temperature treatments during early development (Díaz and Piferrer, 2017). Briefly, E. sea bass larvae were divided into four groups: female control temperature (FCT), female high temperature (FHT), male control temperature (MCT) and male high temperature (MHT) in duplicated tanks. At seven days post fertilization (dpf), FCT and MCT fish remained at 16.5 to 17°C whereas FHT and MHT fish were maintained at 21°C at a ratio of 0.5°C per day. At 68 dpf, all tanks returned to natural temperatures until fish achieved the sex differentiation. At 400 dpf, fish were sacrificed by an overdose of benzocaine and biometric data were obtained. Gonads were dissected and quickly flash frozen in liquid nitrogen and kept at −80°C. For the present experiment, a total of 14 fish were selected: N= 8 males (four fish per treatment), N= 6 females (three fish per treatment). In order to study the heat effects uniquely, the selected fish to further study did not present biometric differences between study groups within the same sex; FCT: 151 g ± 25 g, 21 cm ± 1.29 cm; FHT: 169 g ± 28 g, 22 cm ± 0.95 cm; MCT 77 g ± 17.2 g, 17 cm ± 1.19 cm; MHT: 79 g ± 9 g, 17 cm ± 1 cm, *p-*values > 0.05).

### miRNA extraction

miRNAs of 14 gonads (eight testes and six ovaries) were obtained by miRNAs isolation commercial kit (Qiagen® miRNA, 217004). Quality of the samples was assessed by BioAnalyzer (2100 Bioanalyzer, Agilent Technologies) with ratio 260/280= 2.07 ± 0.04 and 260/230=1,73 ± 0,24) and by the RNA Integrative Number (RIN) measured with a Bioanalyzer (Agilent Technologies, USA) with values > 9.2 in testis. In ovary, RIN numbers were not considered due to naturally high levels of 5s rRNA and tRNA (Bir et al., 2023; Mazabraud et al., 1975)

### miRNA sequencing

In total, 14 libraries were constructed from E. sea bass gonad samples. Library preparation was performed by NEBNext® Small RNA Library Prep Set for Illumina® (Multiplex Compatible) kit following manufacture instructions, using sequencing Lane (1×50, v4, HiSeq) single-end mode with a read length of 50 bp at the Genomics Unit of the Centre for Genomic Regulation (CRG) service in Barcelona.

### Bioinformatics and statistical analysis

Alignment was done using Prost! (Desvignes et al., 2019) using the *Dicentrarchus labrax* genome assembly (version: dlabrax2021) and *Oryzias latipes* was used as reference genome for annotating the miRNAs. miRNAs without annotation were searched in the Fishmirna.org database and complemented (Desvignes et al., 2022). Raw reads were normalized using the DESeq2 package in R and hierarchical clustering of the sample groups was determined by a Principal Component Analysis (PCA) using the plotPCA library in the DESeq2 package (Love et al., 2014). Differential expression was determined by the DESeq2 package in R software. Significant differentially expressed miRNAs (DE-miRNAs) were identified using an *adjusted p*-value (*adj p*-value) cutoff of <0.05. Visualization of differential expression was accomplished by a heatmap using the pheatmap package (version 1.0.12) in R. Target genes of DEM were determined using TargetScan (v. 6.0).

### Quantitative PCR validation

Validation of the miRNA sequencing data was done by quantitative PCR (qPCR) of twelve selected sequenced miRNAs in all the samples used by RNA-seq. cDNA was generated using the miRNA 1st-Strand cDNA Synthesis Kit (Agilent Technologies) following manufacturer’s instructions. Firstly, the polyadenylation reaction was performed after cDNA synthesis. qPCR was performed using the qPCRBio SyGreen blue mix low ROX (PCR Biosystems). A mix of 5 µL 2x qPCRBIO SyGreen Blue mix, 0.4 µL forward primer, 0.4 µL universal reverse primer (Agilent Technologies), 100 ng cDNA and H_2_O up to 10 µL was made for each sample. The sequences of the forward primers for the selected miRNAs were as follows: ola-miR-146a-3p: ATCTATGGGCTCAGTTCTTTTG, ola-miR-7132b-3p: TGAGGCGTTTAGAACAAGTTCA, ola-miR-143-3p: TGAGATGAAGCACTGTAGCTC, ola-miR-21-5p: TAGCTTATCAGACTGGTGTTGG, ola-miR-26a-2/3-b-5p: TTCAAGTAATCCAGGATAGGCT, ola-miR-199a-5p: CCCAGTGTTCAGACTACCTGTTC, ola-miR-199a-3p: ACAGTAGTCTGCACATTGGTT, ola-223-3p: TGTCAGTTTGTCAAATACCCCA, ola-192-5p: ATGACCTATGAATTGACAGCC, ola-miR-194a-5p: TGTAACAGCAACTCCATGTGGA, ola-miR-726-5p: GGAATTCCGCTAGTTCTGAACT, ola-miR-726-3p: TTCACTACTAGCAGAACTCGG. The small nuclear RNA dre-U6: ACTAAAATTGGAACGATACAGAGA was used as the reference gene. A total of twelve comparisons for validations were performed as follows: ola-miR-146a-3p: ovary high temperature (FHT) *vs.* ovary control temperature (FCT), ola-miR-7132b-3p: FHT *vs*. FCT, ola-miR-143-3p: FHT *vs*. FCT, ola-miR-21-5p: testis high temperature (MHT) *vs*. testis control temperature (MCT), ola-miR-26a-2/3-b-5p: MHT *vs*. MCT, ola-miR-199a-5p: MHT *vs*. MCT, ola-miR-199a-3p: MHT *vs*. MCT, ola-223-3p: MHT *vs*. MCT, ola-192-5p: MHT *vs*. MCT, ola-miR-194a-5p:MTHT *vs*. MCT, ola-miR-726-5p: FCT *vs*. MCT, ola-miR-726-3p: FCT *vs*. MCT.

### Conservation analysis

Expression data from miRNAs in ovary and testis were downloaded from the nine fish species in which gonadal miRNAs data were available from Fishmirna.org (Desvignes et al., 2022): zebrafish (Desvignes et al., 2014), three-spined stickleback (*Gasterosteus aculeatus*) (Desvignes et al., 2019), bowfin (*Amia calva*) (Pasquier et al., 2016), striped catfish (*Pangasianodon hypophthalmus*) (NCBI SRA: PRJNA256963), Japanese medaka (*Oryzias latipes*) (Gay et al., 2018), spotted gar (*Lepisosteus oculatus*) (Braasch et al., 2016), black bullhead (*Ameiurus melas*) (NCBI SRA: PRJNA730692), European perch (*Perca fluviatilis*) (Pasquier et al., 2016) and eastern mudminnow (*Umbra pygmaea*) (Pasquier et al., 2016).

Seed region reads from all the gathered datasets were filtered using a threshold of ≥1 count per million (CPM). Common miRNA seed regions between species were visualized in an upset plot using the UpSetR package (version 1.4.0). The top conserved miRNAs were visualized in a dotplot using ggplot2 package (version 3.4.2). Sex-specific overexpression was determined by calculating testis seed region normalized read count divided by ovary seed region normalized read count internally per species. When a seed region was not expressed in ovary or testis, a value of 1 was assigned to avoid division by 0. Following that, the log2 ratio was calculated for all values. To identify highly expressed miRNAs in the ten studied fish species, the relative abundance was shown by calculating the percentage of miRNA abundance in the whole miRNome of the tissue, either ovary or testis. The conserved expression along evolution was determined using a threshold of log2ratio≥|1| in at least 5 species, among which *D. labrax* was mandatory. To show evolutionary relation between the teleost species used in the present study, a phylogenetic tree was generated using PhyloT (v2) website tool.

## Results

### miRNA sequencing and annotation

On average, 10.3 and 6.7 million sequences per library were obtained in testes and ovaries, respectively. The total number of sequences exceeded 146 million; ∼93 and ∼54 million for testes and ovaries, respectively. A total of 299 mature miRNAs were identified after the alignment to the *Dicentrarchus labrax* genome (version dlabrax2021) and after performing the annotation by using the *Oryzias latipes* genome (version ASM223467v1) (Dataset 1). Ten mature miRNAs were not aligned to the genome but were identified using Fishmirna.org (Desvignes et al., 2022) and manually annotated, thus, 100% of the obtained sequences were annotated. Raw sequencing data generated during the current study was submitted in NCBI SRA repository with the accession number: PRJNA1008584. The raw data will be publicly available upon publication and the metadata can be accessed by reviewers through: https://dataview.ncbi.nlm.nih.gov/object/PRJNA1008584?reviewer=ct6r6cbgav3c2ru7evoqvlkqg7.

### Sexual dimorphisms in the miRNAs expression

When comparing testis *vs*. ovary, 69 miRNAs were found to be DE in E. sea bass (*adj p-value*<0.05), of which 33 were upregulated in ovary and 36 were upregulated in testis (Figure 1, Dataset 2). The five most upregulated miRNAs in ovary were miR-734-3p, miR-130c-1-5p, miR-187-3p, miR-155-5p and miR-9-4-3p while in testis were miR-726-3p, miR-724-5p, miR-135b-5p, miR-184a-3p, and miR-726-5p.

**Figure 1.**
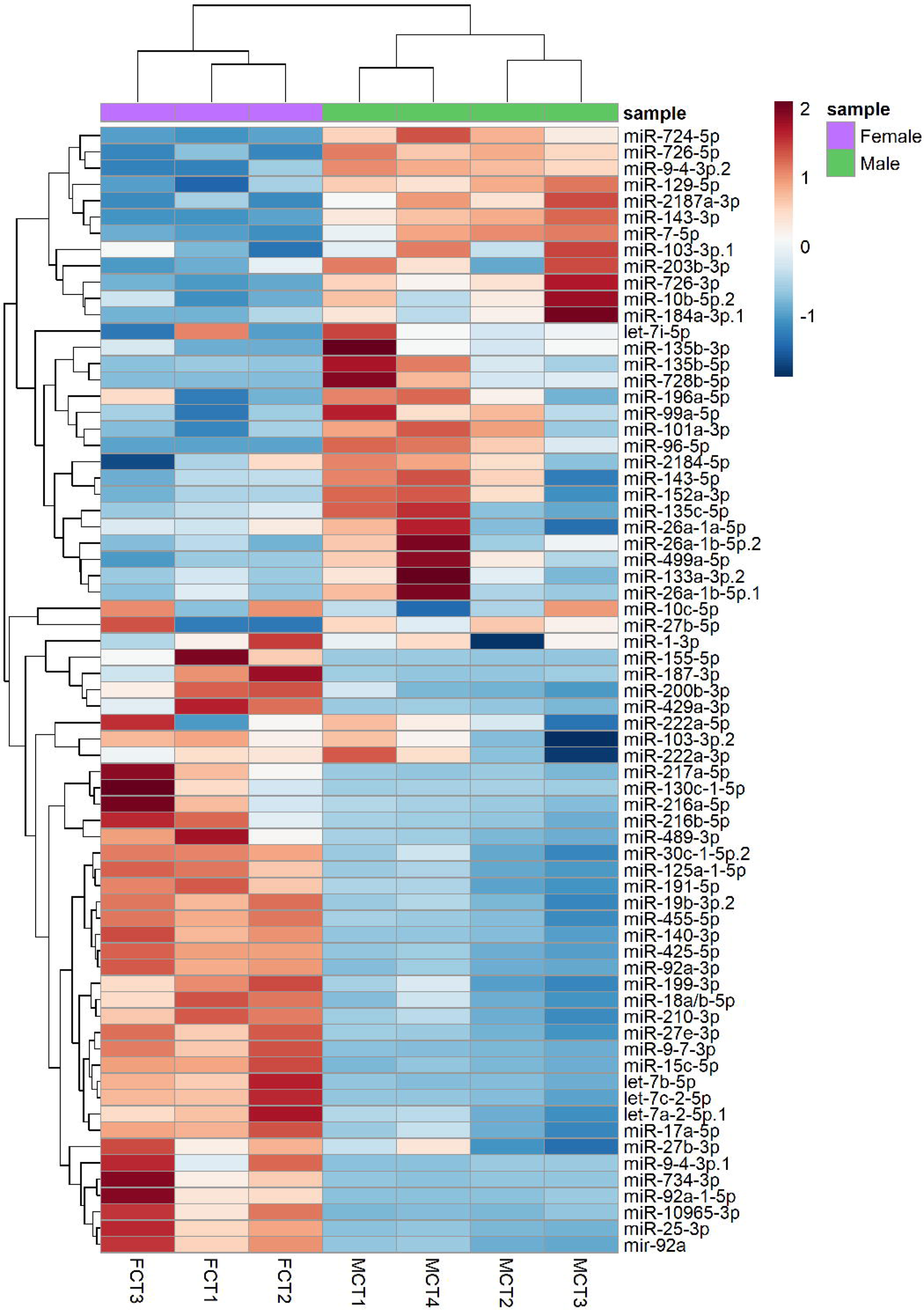
Heatmap of 69 differentially expressed miRNAs (DEM) in European sea bass testes in comparison to ovaries (MCT vs. FCT). The color scale ranges from red to blue, where red shows relative upregulation in testis and blue shows relative upregulation in ovary. Both DEM and samples (N=14) were grouped by hierarchical clustering.

### Heat sensitive miRNAs

FHT vs. FCT and MHT vs. MCT miRNA expression clustering was shown in a PCA (Suppl Figure 1). The two components together explained 57% and 69% of the variance in ovary and testis, respectively.

Almost one year after the end of the heat exposure period development, three miRNAs remained DE (*adj p*-value<0.05) in ovary, one upregulated, miR-192-5p, and two downregulated, miR-143-3p and miR-146a-3p (Dataset 2). In testis, twelve DE miRNAs (DEM) were identified (*adj p*-value<0.05, Figure 2, Dataset 2), of which four (e.g., miR-194a and miR-223-3p) and eight (e.g., miR-92a-a-5p, miR-726-3p, miR-724-5p, miR-143-3p, miR-129-5p) miRNAs were upregulated and downregulated, respectively. Reproduction-related genes were identified as target genes of male DEM, such as *tdrd6* (miR-1388-5p, miR-194a-5p, miR-2187a-3p, miR-9-4-3p and miR-92a-1-5p), *sox3* (miR-129-5p, miR-223-3p, miR-7133-3p), *pgrmc1* (miR-365-2b-3p, miR-7133-1-3p, miR-724-5p) and *gper1* (miR-129-5p, miR-1388-5p, miR-2187a-3p, miR-7133-1-3p, miR-724-5p, miR-9-4-3p, miR-92a-1-5p).

**Figure 2.**
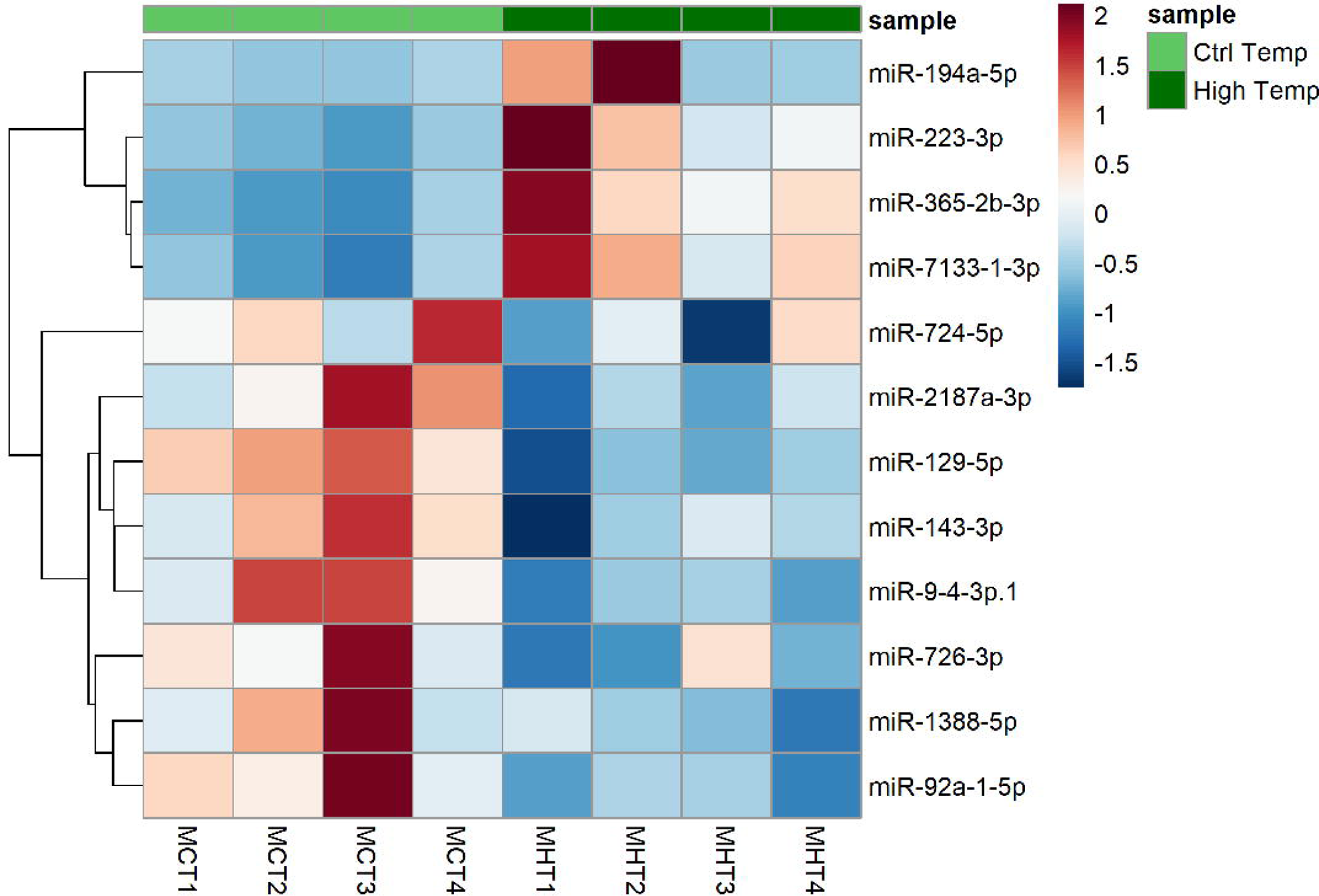
Heatmap of 12 differentially expressed miRNAs (DEM) in European sea bass differentiated testes one year after heat exposure. The color scale ranges from red to blue, where red shows relative upregulation in male high temperature (MHT) males and blue is relative upregulation in male control temperature (MCT).

### miRNA conservation

In order to determine conserved expression along evolution in many fish species, miRNA gonadal data from nine fish species were used together with that of E. sea bass of the present study. By using an upset plot, data showed three intersecting sets (Figure 3): 1) the number of common seed region sequences between the ten species; 2) those seeds that were conserved in less than ten species and 3) the miRNA seeds per each species. In total, 210 seed regions belonging to 105 miRNAs in ovaries (Figure 3A) and in testes 184 seed region sequences belonging to 98 miRNAs were expressed in all ten species (Figure 3B). The evolutionary distances between species is shown in a phylogenetic tree in Suppl figure 2 in showing two clades of the studied species Holostei and Teleostei, emerged ∼250 and ∼310 million of year, respectively.

**Figure 3.**
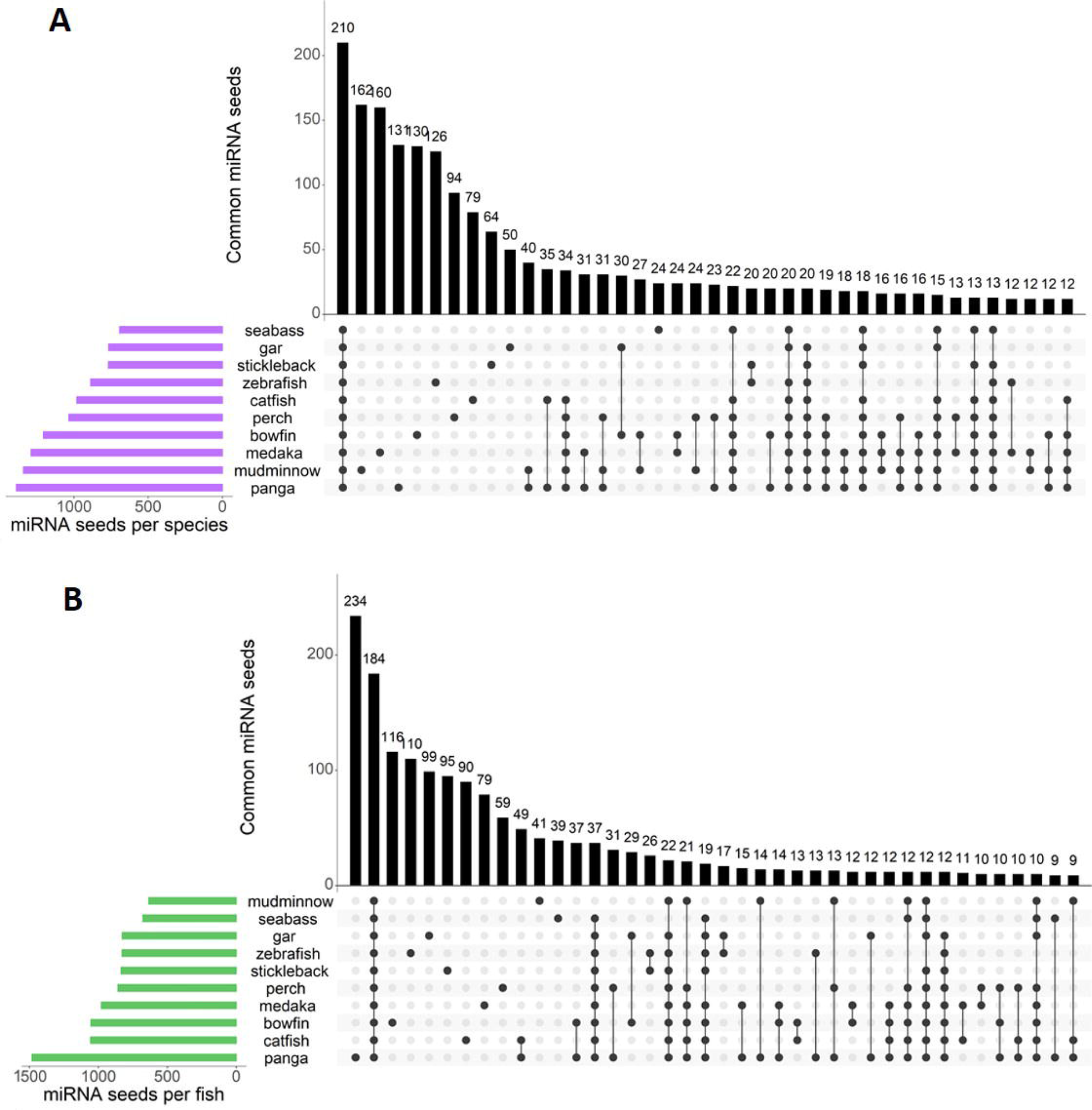
Upset plot of conserved seed regions in ovary (A) and testis (B) in ten teleost species analyzed: *Danio rerio*, *Dicentrarchus labrax, Gasterosteus aculeatus*, *Amia calva*, *Pangasianodon hypophthalmus*, *Oryzias latipes*, *Lepisosteus oculatus*, *Ameiurus melas, Perca fluviatilis* and *Umbra pygmaea*. In ovary (green), 210 seed region sequences were conserved in all ten species. In testis (purple), 184 seed region sequences were conserved in all ten species.

### Abundancy of gonadal miRNAs expression along evolution

To identify highly expressed miRNAs expression patterns throughout evolution in fish, the relative abundance of the top 10 most expressed miRNAs was calculated (Figure 4). In ovary, five miRNAs were highly conserved: miR-143-3p, miR-26a-5p, let-7a-5p miR-100-5p and miR-30a-5p in all ten species (Figure 4A) whereas in testis, two miRNAs, namely miR-143-3p and miR-26a-5p, were identified in all ten species (Figure 4B). The ten highest expressed miRNAs per species in ovary and testis can be found in Dataset 3.

**Figure 4.**
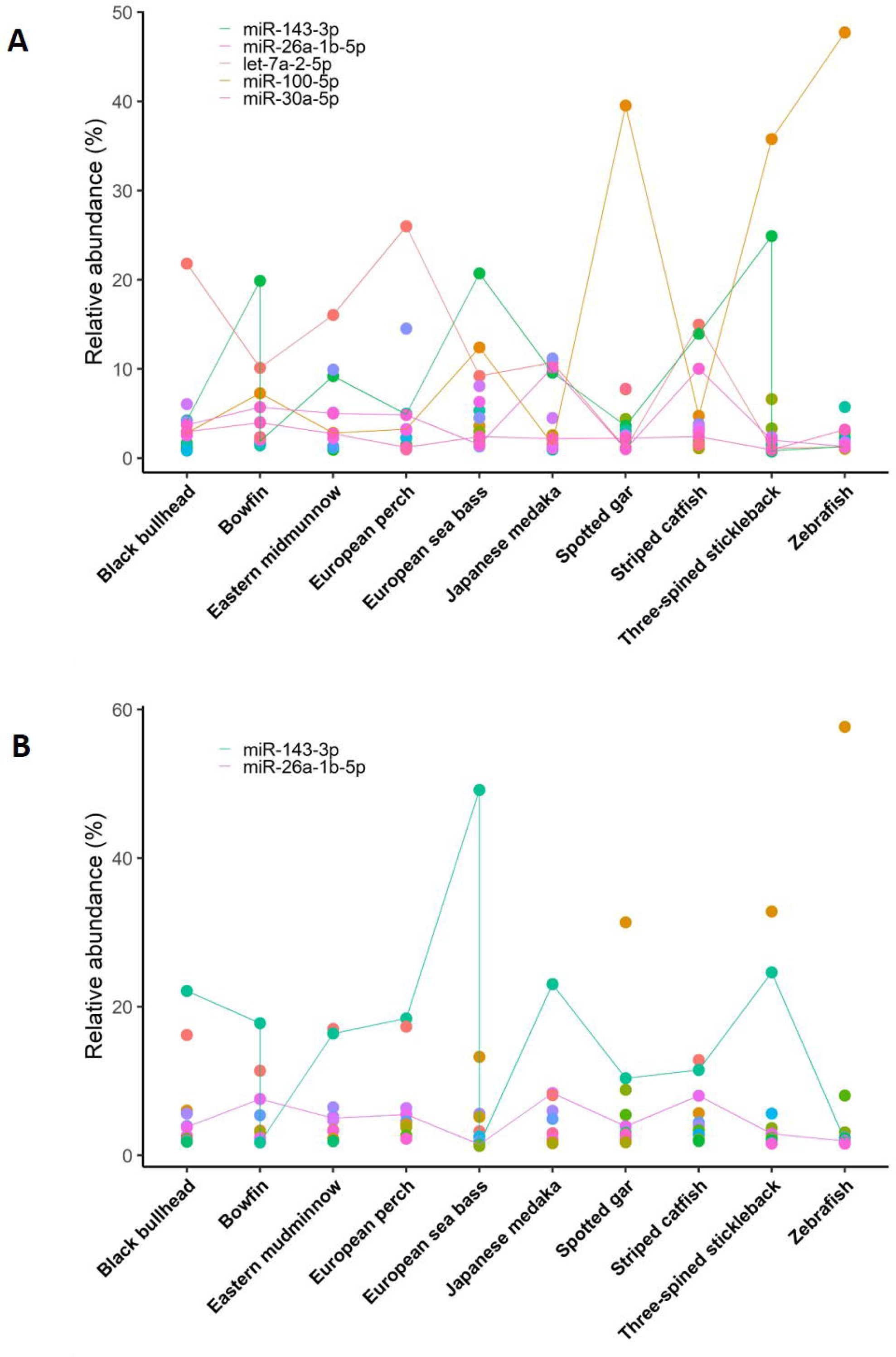
Dot plot of the top 10 most abundant miRNAs in ovary (A) or testis (B) of ten different teleost species. miRNAs expressed in all ten species are shown by a continuous line. The other miRNAs that were in the top 10 most abundant miRNAs in the ovary or testis were plotted by dots with different colors. The top 10 miRNAs per species in ovary and testis can be found in Dataset 3.

### miRNA gonadal markers

To identify sex-skewed miRNA expression in different fish species and miRNAs markers as heat recorders, the following analyses were performed. First, miRNAs were identified with a log2 ratio>|1| of testis *vs*. ovary in at least five or more species resulting a total of 83 miRNAs. From those 49 identified as female-biased miRNA markers and 34 as male-biased markers because they were more expressed either in the ovaries or testes in at least five species along evolution (Suppl Table 1). Secondly, miRNAs were further filtered based on the heat-treated data obtained in E. sea bass in the different comparisons based on sex and heat treatment (Table 1): (1) MHT *vs.* MCT, (2) FHT *vs.* FCT, and (3) MCT *vs*. FCT. This resulted in a total of 19 miRNAs, eleven of which were female-like miRNA markers, for example, miR-10965-3p, miR-155-5p, miR-223-3p and miR-425-5p and eight as male-like miRNA markers, for example, miR-2187a-3p and miR-143-3p. Most of the species showed a similar sex-skewed patterns of these miRNA markers, except *G. aculeatus* and *A. melas* species, which their expression patterns differed, in particular, for those identified as female-like miRNA markers.

**Table 1.** Sex-skewed miRNA expression in different fish species and the effects of heat treatment. miRNAs were identified with log2 ratio>|1| of seed region expression in testis *vs.* ovary in at least five or more fish species, among which E. sea bass. Furthermore, each miRNA was significantly differential expressed (DEM, *adjusted-p* value<0.05) in our data. The E. sea bass DEM comparisons were: (1) Male high temperature (MHT) *vs.* Male control temperature (MCT); (2) Female high temperature (FHT) *vs.* Female control temperature (FCT); (3) MCT *vs*. FCT. Negative values mean the upregulation in ovary whereas positive values mean upregulation in testesfor each species. The miRNA name corresponds to the seed region annotation aligned with European sea bass (*Dincentrarcus labrax*). Data in the table was organized based on the miRNA conservation in the ovaries or testes along fish evolution. Abbreviations: dla: *Dicentrarchus labrax*; ola: *Oryzias latipes*; aca: *Amia calva*; gac: *Gasterosteus aculeatus*; dre: *Danio rerio*; loc: *Lepisosteus oculatus*; phy: *Pangasianodon hypophthalmus*; pfl: *Perca fluviatilis*; ame: *Ameiurus melas*; upy: *Umbra pygmaea*

| Seed | miRNA | dla | ola | aca | gac | dre | loc | phy | pfl | ame | upy | Sea bass DEM |
| --- | --- | --- | --- | --- | --- | --- | --- | --- | --- | --- | --- | --- |
| AAAGAGG | miR-10965-3p | -5.12 | -0.12 | -5.42 | 0.00 | -3.05 | 0.00 | 0.00 | -7.21 | -4.32 | -4.36 | 3 |
| TAATGCT | miR-155-5p | -4.15 | -3.80 | -1.31 | 0.47 | -0.36 | -0.66 | -1.64 | -4.26 | -0.18 | -0.45 | 3 |
| ATACTGT | miR-429a-3p | -3.33 | -2.68 | -3.22 | 2.44 | 0.00 | 0.00 | -0.33 | -6.04 | 1.77 | -4.26 | 3 |
| ATGACAC | miR-425-5p | -2.88 | -2.01 | -0.86 | -0.21 | 0.61 | -1.12 | -1.31 | -2.81 | -0.03 | 0.08 | 3 |
| GTAACAG | miR-194a-5p | -2.27 | -1.39 | -0.80 | 3.99 | -1.42 | -0.55 | -3.16 | 2.23 | 4.23 | -2.23 | 2 |
| TGACCTA | miR-192-5p | -2.06 | -1.47 | -1.31 | 4.91 | -0.62 | -0.27 | -2.97 | 2.25 | 3.67 | -2.46 | 2 |
| AACGGAA | miR-191-5p | -2.03 | -2.08 | -0.06 | 1.12 | 0.79 | -1.18 | -1.30 | -1.88 | -0.16 | 0.02 | 3 |
| TGACATC | miR-489-3p | -1.90 | -2.21 | 0.00 | 0.29 | -1.74 | -1.14 | 2.27 | -1.87 | 5.20 | 0.00 | 3 |
| ACCACAG | miR-140-3p | -1.65 | -0.97 | -1.01 | 1.56 | -0.53 | -2.28 | 0.01 | -1.64 | 1.15 | -2.37 | 3 |
| ATACTGC | miR-200b-3p | -1.35 | -3.20 | -3.16 | 3.54 | -2.75 | -3.51 | 0.82 | -0.80 | 2.04 | -1.64 | 3 |
| GTCAGTT | miR-223-3p | -1.18 | -4.17 | -1.13 | -0.19 | -1.83 | 0.48 | -0.13 | -1.52 | 0.57 | 0.34 | 2 |
| GAGATGA | miR-143-3p | 1.25 | 1.26 | -0.16 | -0.02 | 0.63 | 1.52 | -0.28 | 1.91 | 2.39 | 0.83 | 1+2+3 |
| GGACGGA | miR-184a-3p | 2.01 | -1.67 | 1.20 | 1.08 | 4.77 | 1.36 | -3.12 | -1.45 | -1.35 | 0.00 | 3 |
| TAAGACT | miR-499a-5p | 2.23 | -3.28 | -1.00 | 1.29 | 3.94 | 1.94 | -1.95 | 2.82 | 2.27 | 3.71 | 3 |
| TTTTTGC | miR-129-5p | 2.38 | -0.03 | 3.44 | -3.80 | -0.30 | 1.46 | -0.75 | 1.68 | 4.14 | 6.31 | 1+3 |
| TTACAGG | miR-2187a-3p | 2.51 | 2.43 | 1.54 | 0.00 | 1.65 | 4.07 | -0.41 | 0.00 | 3.30 | 0.00 | 1+3 |
| GGAAGAC | miR-7-5p | 2.83 | 4.37 | -1.26 | -3.05 | 0.80 | 1.43 | -2.11 | 1.83 | 2.66 | 0.83 | 3 |
| TAAAGGG | miR-724-5p | 3.84 | -0.07 | -1.97 | 5.48 | -3.62 | 2.02 | 1.17 | 0.89 | -3.06 | 4.90 | 1+3 |
| TATAGGG | miR-135b-3p | 4.62 | 2.24 | 0.28 | 3.77 | 0.00 | -1.20 | -0.07 | 0.00 | 4.20 | 5.93 | 3 |

Noteworthy, from the comparison (1), a total of six miRNAs altered the expression after one year of the heat treatment in males. Four of them were downregulated in testes subjected to HT in E. sea bass but also identified as male-like miRNAs, which were miR-129-5p, miR-2187a-3p, miR-724-5p and miR-143-3p (Figure 5). The other were two upregulated miRNAs, i.e., miR-223-3p and miR-194a-5p, which were identified as being upregulated in ovaries along evolution, therefore as female-like markers. Further, for the comparison (2), two miRNAs were altered in the heat-treated ovaries, miR-192 which was upregulated and identified as female-like marker and miR-143 which was downregulated and identified as male-like marker. The miR-143-3p was identified as DEM in all the three comparisons studied (1), (2) and (3), indicating that this miRNA was highly conserved in testes along evolution and it was heat sensitive in E. sea bass, thus showing in our analysis, a downregulation in both heat-treated testes and ovaries (Figure 5).

**Figure 5.**
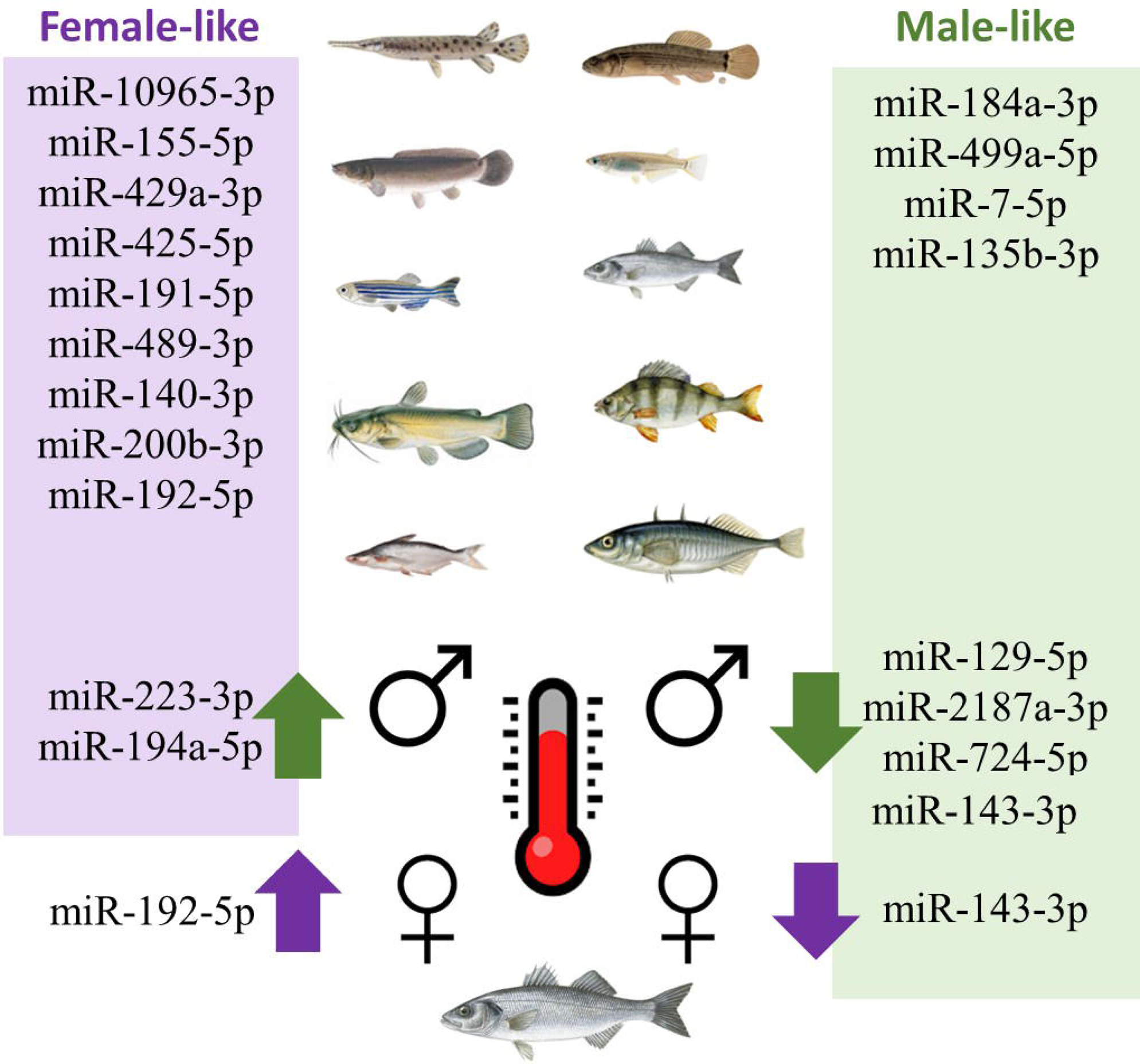
Gonadal miRNAs markers identified in this study. In purple boxes, the female-like miRNAs were represented and in green, the male-like miRNAs with log2 ratio>|1| of seed region expression in testis *vs.* ovary in at least five or more fish species. Heat-recorder miRNAs were significantly differential expressed (DEM, *adjusted-p* value<0.05) in the European sea bass after one year of heat treatment during early development.

### RNA-seq validation

To validate the RNA-seq data, qPCR analysis was performed on the ovaries and testes samples. When comparing the log2FC as obtained by qPCR and RNA-seq for twelve miRNAs, we obtained an R^2^=0.9426 and *p*-value=1.6e-07 (Suppl Figure 2) thus validating our data.

## Discussion

High temperatures are recognized to induce masculinization of fish populations. This phenomenon has significant implications for many farmed species, including the E. sea bass, in which females grow to a larger size than males. In addition, increasing temperatures in our seas and oceans due to climate change has drastic implications in fish physiology and evolution (Alfonso et al 2020). Here, we study the effect on the gonadal miRNome of this heat-induced masculinization and further explore the gonadal miRNome of other species to decipher miRNA sex markers.

Long-term effects of heat treatment on the miRNA expression profile in the gonads has been previously described in fish. In particular, in Atlantic cod, miR-27c, miR-30c, and miR-200a were found to be DE in juveniles after embryonic or larval heat-treatment, however, no distinction between males or females was addressed (Bizuayehu et al., 2015). Other studies in zebrafish describing the heat stress by temperature in the gonadal miRNome revealed that miR-22b and miR-301a were altered to be fully recovered three months after the exposure (Ikert and Craig, 2020). In zebrafish females, 23 miRNAs were DE in ovaries after larval heat-treatment, among which, for example, dre-miR-726-3p was downregulated while dre-miR-724-5p was upregulated (van Gelderen et al., 2022). Interestingly, these two miRNAs were both found to be downregulated in E. sea bass testis one year after heat exposure in the present study, indicating that a conservation in the heat response among species may exist. Similarly, miR-143-3p and miR-92a-3p, which were downregulated in high temperature treated fish in the present study, were altered in mice testes exposed for only 25 minutes to elevated temperatures, highlighting them as potential markers in heat-induced spermatogenesis disorder (Gan et al., 2023). Higher levels of circulating miR-143 and miR-223 were detected in plasma levels together with cortisol after acute stress response in E. sea bass (Houdelet et al., 2023), similarly of what observed in this study by heat stress. Overall data shows that in fish, miRNAs play a regulatory role in the thermal plasticity of acclimation and adaptation to drastic temperature changes in the environment.

In E. sea bass, we observed sexual dimorphism in miRNA expression within fish gonads, with 33 upregulated and 36 downregulated miRNAs in the ovary compared to the testis. To enhance reliability, we cross-referenced our data with existing datasets from the nine species, identifying common gonadal miRNAs along evolution. In testes and ovaries, we have identified 184 and 210 conserved seed region sequences in the ten studied fish species, respectively. Although we have detected the expression of these miRNAs in the gonads in many fish species, these miRNA seed sequences were not sex-specific or tissue-specific, as many miRNAs are highly conserved throughout evolution and body plan development. For example, miR-100 and let-7, were considered as an “ancestral miRNAs”, due to its conservation already described within Metazoan species (Christodoulou et al., 2010; Grimson et al., 2008; Hertel et al., 2012). Nevertheless, a conservation analysis revealed 74 conserved miRNAs divided into three groups: 23 miRNA families were present in both protostomes and deuterostomes, 46 families conserved exclusively in vertebrates, and five families (mir-430, mir-722, mir-724, mir-734, and mir-738) exclusively for fish species (Li et al., 2014). Notably, in our data, miR-143-3p, miR-26a-5p, let-7a-5p, miR-100-5p and miR-30a-5p in the ovaries, and miR-143-3p and miR-26a-5p in the testes were the most expressed miRNAs in all ten studied fish species. Similarly, in olive flounder (*Paralichthys olivaceus*), miR-143, miR-26a and let-7a were also identified as the most expressed miRNAs in both ovaries and testes (Gu et al., 2014). In mouse and human, miR-30a-5p was crucial for spermatogonial stem cell differentiation (Khanehzad et al., 2021) and was used as a marker for men suffering from non-obstructive azoospermia (Arefnia et al., 2022). In our data, however, we observed a conserved overexpression of miR-30a in adult fish ovaries compared to adult testes in the ten studied species. Nevertheless, in Nile tilapia, miR-30a presented higher expression in males at 5 days after hatching (dah) and it was able to downregulate *cyp19a1a* in gonads (Tao et al., 2016). We identified both in ovaries and testes the miR-26a, which in Chinese tongue sole (*Cynoglossus semilaevis*), was identified as transgenerational male marker in sperm (Zhao et al., 2021). Overall, data show the complexity of the functions of miRNA along evolution and their multiple role in regulating many biological processes even those involved in the reproduction system.

In fact, the exploration of miRNA phylogenetic conservation and diversity indicates that miRNAs play crucial roles in animal evolution by influencing phenotypic variation during development (Niwa and Slack, 2007b). This might explain the different sexual patterns observed in the gonads between the ten fish species of two different clades (i.e., Teleostei and Holostei) separated from one species to others for millions of years of evolution. Further, representing distinct reproduction systems, from XX/XY chromosomal system such as medaka, stickleback, perch and, panga (Matsuda et al., 1998; Peichel et al., 2020; Rougeot et al., 2002; Wen et al., 2022), to polygenic system such as E. sea bass and zebrafish (Liew et al., 2012; Vandeputte et al., 2007). Thus, to detect sexual dimorphic miRNA markers in the fish gonads, we considered a sex-biased miRNA when the expression was found in at least five studied species. In total, we identified eleven and eight female-like and male-like markers, respectively. For example, miR-429a-3p and miR-140-3p were overexpressed in ovary in at least five studied species. miR-429b-3p was reported to have lower expression in yellow catfish in YY super males compared to XY males (Jing et al., 2014) and in chicken, miR-140-3p was shown to promote germ cell proliferation while targeting anti-Müllerian hormone *amh* (Pfennig et al., 2015; Zhang et al., 2023). On the other hand, a male-like miRNA such as miR-184a-3p was shown to be a highly expressed miRNA in testis and involved in spermatogenesis in mice (Wu et al., 2011). Scarce data exists about miR-499, which has been related in cardiac disorders, lung cancer and regulating circadian clock (Ahmed et al., 2023; Chen et al., 2013; Pisano et al., 2015) in mammals. In fish, miR-499 plays a role in muscle tissue (Duran et al., 2015; Nachtigall et al., 2015, 2014) and its evolution targeting the intronic region of the myosin heavy chain (MYH14) gene was studied throughout evolution in teleost (Bhuiyan et al., 2013). In a recent publication in E. sea bass, the authors identified higher levels of circulating miR-499 in plasma differentiating males when comparted to differentiating females (Houdelet et al., 2023). Another example of a male-like miRNAs in fish gonads is the miR-135, which is a biomedical marker in humans due to its involvement in many cancer and disorders like Alzheimer disease (Kadkhoda et al., 2022; Zheng et al., 2021). Similar to our data, in tilapia, miR-135 presented sexual dimorphism towards males as it was upregulated in testes together with miR-33a, miR-132 and miR-212 (Herkenhoff et al., 2018; Xiao et al., 2014).

To identify miRNA markers that convey information about both sexual dimorphism and serve as recorders of environmental cues, we considered the sex-bias miRNAs together with the data from the heated E. sea bass. These analyses aided in pinpointing miRNAs that could be accountable for the masculinizing effects induced by high temperatures detecting a total of seven informative miRNAs. miR-194a-5p and miR-223-3p were upregulated in HT males but overexpressed in ovaries in at least five fish species, thus indicating that they might be involved in the masculinization of the gonads and thus, the differentiated testes might belong to genetic females. In Chinese sole tongue (*Cynoglossus semilaevis*), miR-223-3p (referred to as miR-223-y) was upregulated during oocyte maturation and targeted the insulin-like growth factor 1 receptor (*igf1r*), a gene involved in testicular development (Cannarella et al., 2018). The expression of miR-223-3p was lower in YY super males of yellow catfish (*Pelteobagrus fulvidraco*) than XY males (Jing et al., 2014). On the other hand, male-like miRNAs that showed heat-sensitivity were miR-129-5p, miR-2187a-3p, miR-724-5p and miR-143-3p which were downregulated in HT males. In literature, miR-129 was associated with sexual maturity in rainbow trout (*Oncorhynchus mykiss*) testis (Farlora et al., 2015) while miR-724-5p (referred to as miR-724-x), showed upregulation during oocyte maturation although it suppressed *cyp19a1a* (Guiguen et al., 2010; Zhang et al., 2022). In mice (Gan et al., 2023), miR-143-3p was shown to be highly upregulated after heat-induced stress reducing spermatogenesis and thus indicating the thermal sensitivity of this miRNAs in the gonads. In all, the identification of these miRNAs may serve as potential epimarkers for predicting past environmental events based on the sexual fish phenotype.

## Conclusions

To gain a deeper understanding of the intricacies of masculinization induced by heat treatments during early developmental stages in E. sea bass, we detected a total of four and twelve miRNAs in the ovaries and testes, respectively. To explore the sexual dimorphic miRNA patterns in the gonads across different fish species, we integrated our findings with available miRNAs from public databases. This analysis revealed six conserved miRNAs throughout evolution (i.e., miR-194a-5p, miR-223-3p, miR-129-5p, miR-2187a-3p, miR-724-5p, and miR-143-3p) as markers for heat-induced masculinization in fish gonads.

## Ethics

Experimental procedures agreed with the European regulations of animal welfare (ETS N8 123,01/01/91) and were approved by the Ethical Committee of Consejo Superior de Investigaciones Científicas (CSIC) that evaluates projects and procedures in which animals are used for experimentation and other scientific purposes –RD 53/2013 (Spanish Ministry of Science and Innovation). All procedures performed were in accordance with the ethical standards of the institution and followed the European Directive 2010/63 UE. The study was carried out in compliance with the ARRIVE guidelines.

## Acknowledgements

This study was supported by the Spanish Ministry of Science and Innovation grant 2PID2020-113781RB-I00 “MicroMet” and by the Consejo Superior de Investigaciones Científicas (CSIC) grant 02030E004 “Interomics” to LR, by grant PID2019-108888RB-I00 to FP, and with funding from the Spanish government through the ‘Severo Ochoa Centre of Excellence’ accreditation (CEX2019-000928-S). We thank Noeli Díaz for sampling the fish, the lab technician Sílvia Joly for her essential assistance in our team and Gemma Fusté for her assistance in fish facilities.

## Supplementary Information

**Supplementary Table S1.** Ratio of seed region expression in male control temperature (MCT) *vs.* female control temperature (FCT) in ten teleost species. Purple: overexpression with log2ratio>1 in ovary, green: overexpression with log2ratio>1 in testis.

## Supplementary Figures

**Supplementary Figure 1. Principal Component Analysis (PCA) of miRNA expression of gonadal samples (A, ovaries and B, testes) of European sea bass after heat treatments during early development.** Abbreviations: FCT, female control temperature; FHT, female high temperature; MCT, male control temperature; MHT, male high temperature.

**Supplementary Figure 2.** Evolutionary relation between the ten teleost fish species used in the present study generated by PhyloT (v2) website tool. https://phylot.biobyte.de/

**Supplementary Figure 3.** Validation of the miRNA sequencing data by quantitative PCR (qPCR) of twelve selected sequenced miRNAs on the ovaries (N=6) and testes (N=8) samples used in the present study. Table shows the results of the two techniques used by comparing the log2FC as obtained by qPCR and RNA-seq for 12 miRNAs.

## Datasets

**Dataset 1.** Prost! output of European sea bass miRNA annotation and reads obtained after miRNA-sequencing of ovaries and testes treated with high temperature during early stages of development.

**Dataset 2.** Differentially expressed miRNAs in different comparisons of E. sea bass gonads treated with high temperature during early gonadal development. Comparisons: male control temperature (MCT) *vs.* female control temperature (FCT), female high temperature (FHT) *vs*. FCT and, MHT *vs.* MCT.

**Dataset 3.** Top 10 miRNAs in ovary and testis for each fish species.

