## Supplemental figures for "Identification of sex-biased miRNA markers informative of heat-past events"

### Slide 1
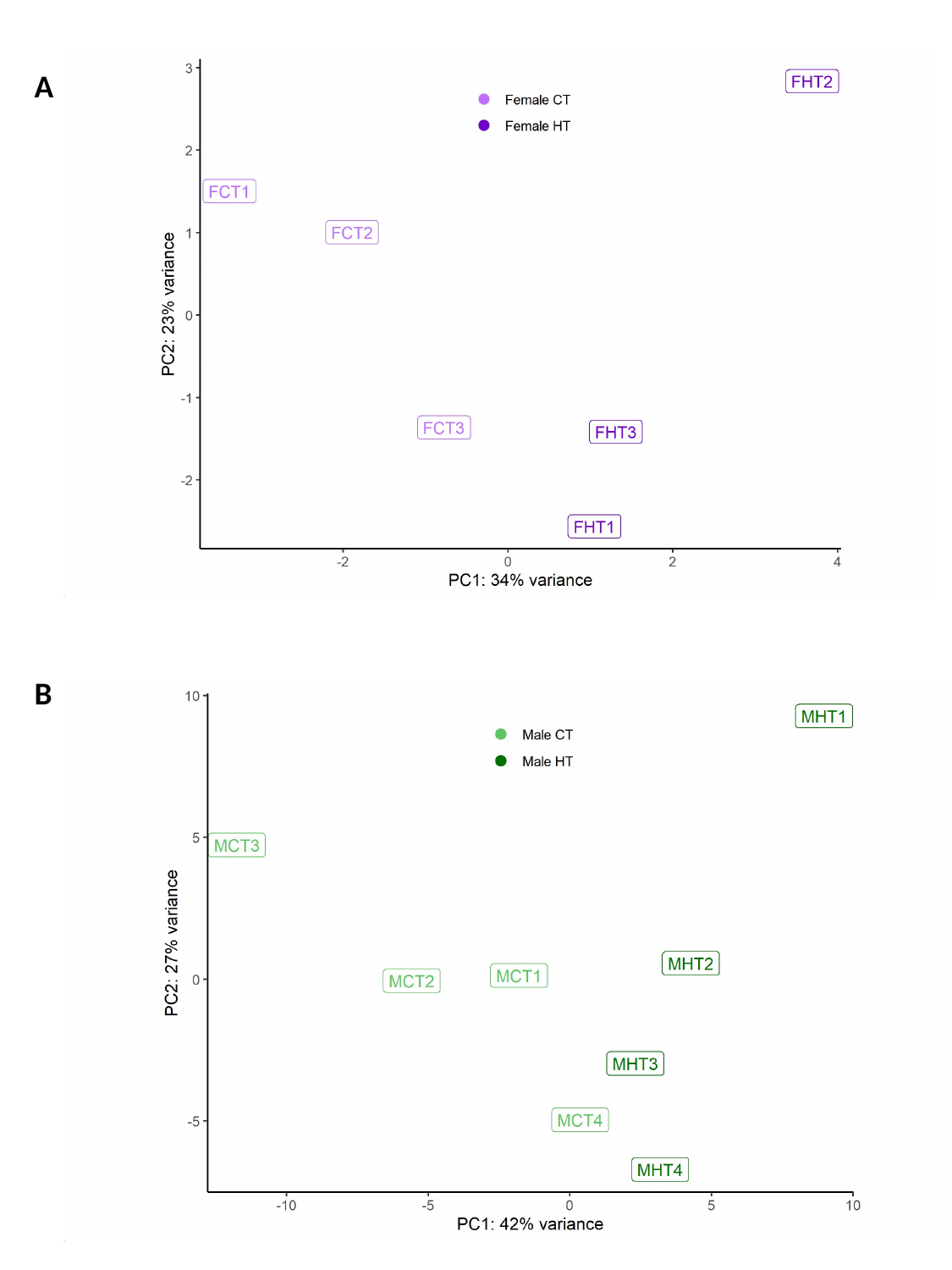

A
B

### Slide 2
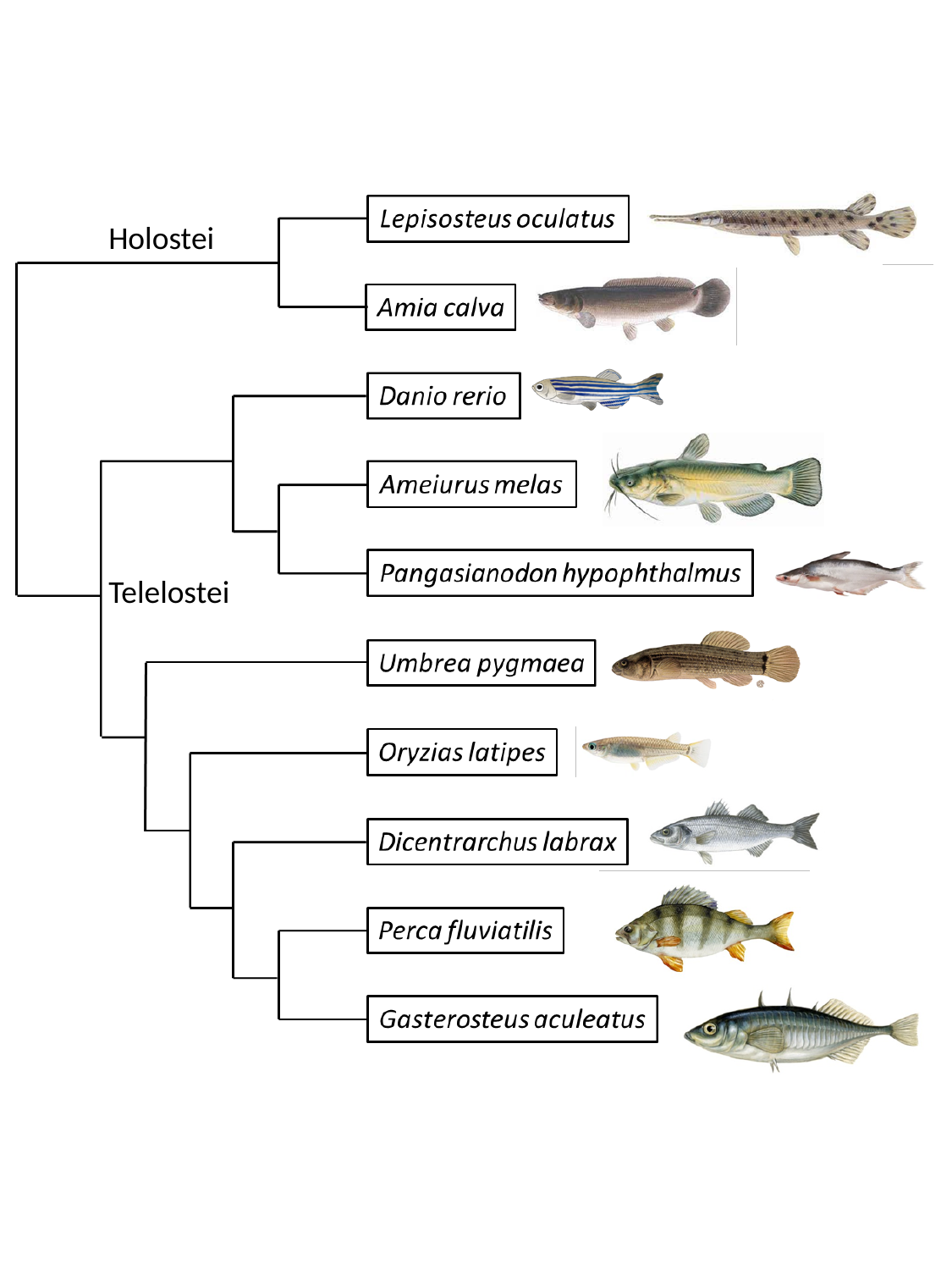

Holostei
Telelostei

### Slide 3
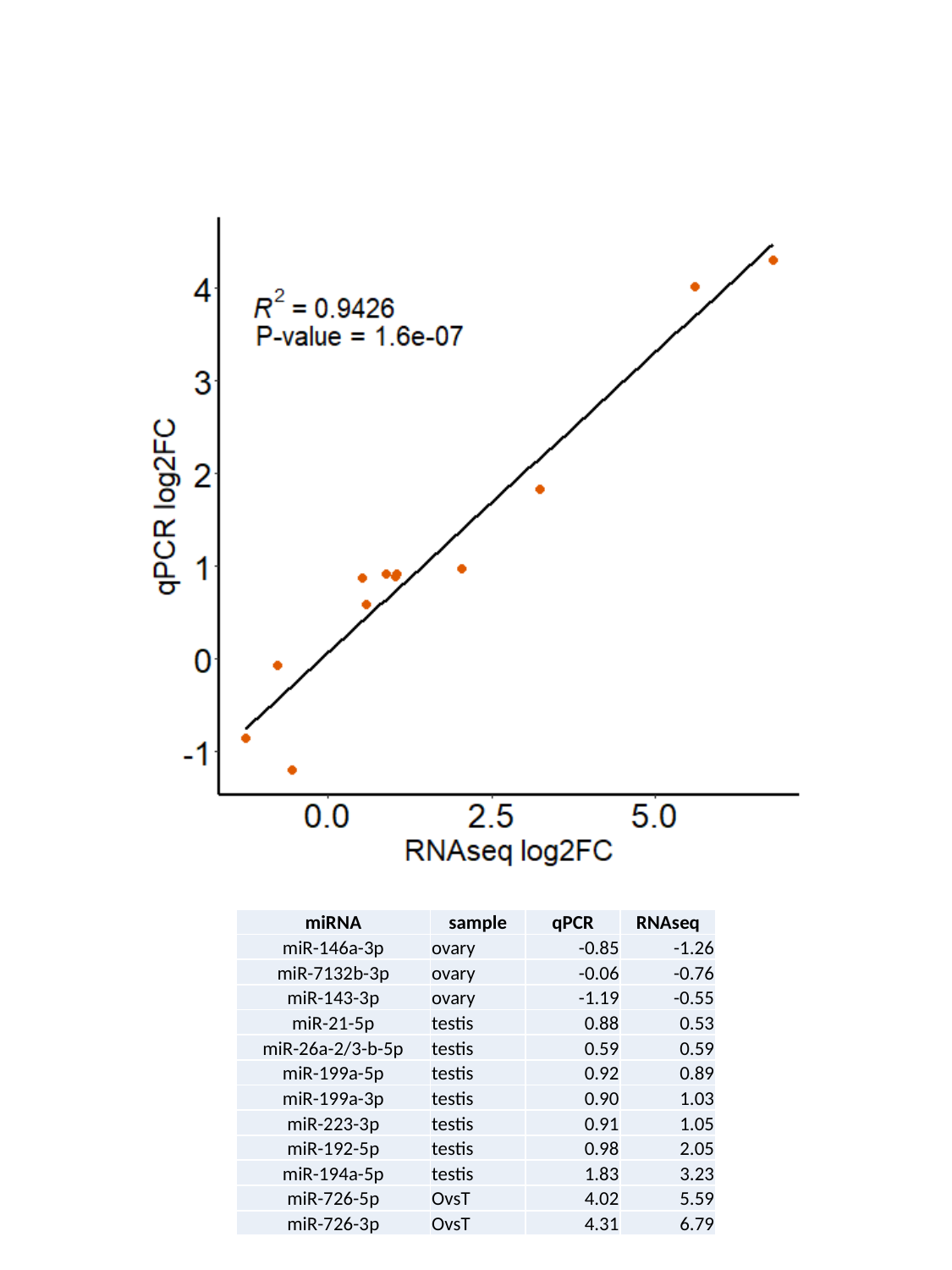

| miRNA | sample | qPCR | RNAseq |
| --- | --- | --- | --- |
| miR-146a-3p | ovary | -0.85 | -1.26 |
| miR-7132b-3p | ovary | -0.06 | -0.76 |
| miR-143-3p | ovary | -1.19 | -0.55 |
| miR-21-5p | testis | 0.88 | 0.53 |
| miR-26a-2/3-b-5p | testis | 0.59 | 0.59 |
| miR-199a-5p | testis | 0.92 | 0.89 |
| miR-199a-3p | testis | 0.90 | 1.03 |
| miR-223-3p | testis | 0.91 | 1.05 |
| miR-192-5p | testis | 0.98 | 2.05 |
| miR-194a-5p | testis | 1.83 | 3.23 |
| miR-726-5p | OvsT | 4.02 | 5.59 |
| miR-726-3p | OvsT | 4.31 | 6.79 |
